# BAG6 is a novel player in controlling nonalcoholic steatohepatitis: result from a comprehensive *in-silico* study

**DOI:** 10.1101/2023.05.04.539506

**Authors:** Dipanka Tanu Sarmah, Abhijit Paul, Umang Berry, Milan Surjit, Nandadulal Bairagi, Samrat Chatterjee

## Abstract

Nonalcoholic steatohepatitis, or NASH, is a multifactorial disease characterized by hepatic lipid accumulation, inflammation, cell death, and fibrosis, and an efficacious pharmaceutical intervention for this is yet to be discovered. The present study aims to identify potential targets capable of reversing the disease-specific molecular alterations and elucidate their possible action mechanism. Our study uses combinations of different methods, such as genome-scale metabolic modelling, directional protein-protein interaction network, connectivity map, and network controllability, to identify potential targets in NASH. Our approach yielded three promising targets, BAG6, CASP3, and CYCS, and captured their effects on inflammation, fibrosis, steatosis, and apoptosis. The association of CASP3 and CYCS with NASH are already reported in the literature. So BAG6 was selected as a novel target. In the Huh-7 cell-line, its ablation reduced fatty acid accumulation and decreased levels of NASH-signature transcripts, supporting our hypothesis on BAG6 as a potential NASH target.

## 1. Introduction

Nonalcoholic fatty liver disease (NAFLD) is a spectrum of liver disease defined by the aggregation of triglycerides in the liver without other causes, such as medications, excessive alcohol intake, or certain heritable conditions [1]. Nonalcoholic steatohepatitis (NASH), characterized by steatosis, lobular inflammation, and hepatocellular ballooning, is the second stage of this continuum. It is a slowly progressive disease that often remains clinically discerned, resulting in late detection, curbing the therapeutic options, and contributing to poor outcomes. Hence, there is a need to reveal the underlying molecular mechanisms of the disease that might lead to the development of effective treatment strategies.

Probing context-specific networks, such as protein-protein interaction (PPI) and/or metabolic networks, is the only way to properly understand the anomalies in the cellular systems that occur with a disease’s progression. Proteins work in conjunction with others to accomplish various cellular processes. The PPI network analysis gives us information about the core set of proteins driving the progression of the disease [2,3]. On the other hand, metabolic networks can investigate alterations in the metabolism that befalls across the entire histological spectrum of disease, probing the changes that develop during the progression of benign to severe stages [4]. Thus, there is a high demand for identifying an ideal target that not only regulates the disease interactome and generates a reverse gene expression profile but is also capable of exerting a significant influence on the altered metabolic pathways under diseased conditions.

Genome-scale metabolic models (GSMMs) capture the altered metabolic pathways through a bottom-up systems-level understanding of the metabolic network. Over the past decade, GSMM has widely been employed to capture the disease-associated molecular mechanisms [5,6], which are further used to identify potential drug targets [6,7] and biomarkers [5]. Some therapeutic strategies are proposed in NAFLD by identifying altered metabolic pathways using GSMM [8,9]. Besides the metabolic alterations, pathways like inflammation, fibrosis, apoptosis, etc., also contribute to the NAFLD progression. Thus, protein and metabolic level rewiring is required to control the disease progression. In this context, protein-protein interaction (PPI) network analysis can be applied to identify a core set of proteins capable of governing the system and hence might be the potential targets that influence all the aforementioned pathways. Among several methods, a reliable perspective for studying such networks is the implications of control theory, which investigates how to manipulate a dynamic system’s behaviour [2,3]. However, most works using the PPI network mainly focus on identifying only the hub genes from the differentially expressed genes (DEG) network constructed from various transcriptomic data [10,11]. The primary limitation of these studies is that they do not consider the changes in metabolic flux level and hence cannot look into the perturbations in vital metabolic pathways driving the disease progression. However, the efficient analysis of the PPI network can provide novel candidates for metabolic channelling. Again, the core set of proteins, which can be the potential drug targets, should be capable of reversing the disease-associated gene signatures. Hence, to better understand the feasibility of the targets, their role in the disease system should be adequately investigated, which brings the collaborative effort of the context-specific molecular networks into the scenario.

Taking all these as motivation, we endeavour to study NASH at protein and metabolic levels to identify potential therapeutic targets. By taking a clinical dataset on NAFLD, the gene and the metabolic level alterations were simultaneously investigated to find the crucial genes that can initiate reversibility towards control. These genes were further integrated into a directional PPI network of the DEGs and applied control theory analysis to get the most fragile nodes in the interactome. The identified proteins can induce reverse gene expression and metabolic transformation towards control and affect network controllability. We then analyzed their knockdown gene expression profile to tailor their effect on steatosis, inflammation, fibrosis, and cell death, the four clinical traits of NAFLD. We looked for novel targets not previously reported to be associated with NASH and evaluated the functional significance of these targets using a cell-based model.

## 2. Materials and methods

### 2.1) *In-silico* gene knockdown exercise using GSMM

We performed an *in-silico* single gene knockdown approach to obtain the gene set responsible for the network perturbations towards the healthy state. To get the disease-specific GSMM, we integrated the average gene expression values into the *iHepatocytes2322* [8] by applying the E-Flux method [12]. On the other hand, the knockdown model was built by considering the 90% reduction in the expression value of a particular gene, and the corresponding flux state (V^res^) was obtained by using minimization of metabolic adjustment (MOMA) [13]. This process requires the flux state (V^ref^) of disease-specific GSMM. The ‘*gpSampler’* function available in the CobraToolbox 3.0 [14] was used for flux sampling, and the mean value of the flux distributions was considered. Finally, we assigned scores for each gene knockdown similar to the method proposed in the metabolic transformation algorithm (MTA) [15], following the rule

Transformation Score 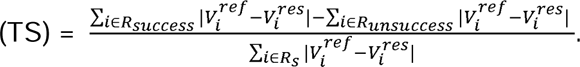.

The altered relations are classified into two groups R_success_ and R_unsuccess_ based on the changes in flux rates in the required direction and R_s_ represents the unaltered reactions.

### 2.2) Identifying candidate proteins based on reversible gene signature patterns using the Connectivity Map database (CMap)

The query tool of the CMap database [16] takes the list of up-and-downregulated genes and provides a connectivity score (ranges from −100 to +100) to perturbagens, mainly based on the similarity between the query gene set and the reference gene set. The higher the positive score, the higher the correlation between the query set and the reference set of genes. Similarly, a negative score means that the induction of that particular perturbagen causes an opposite gene expression profile to the query gene set. For this purpose, we selected the gene whose knockdown has a connectivity score ≤-90.

### 2.3) Gene knockdown profile extraction using the CMap database

we extracted the LINCS L1000 dataset from the CMap database to pip into the effects of the indispensable candidate protein (ICp) in altering the metabolic processes. Here, we have opted for the level 5 dataset in our study as it is more robust [17]. It contains the consensus replicate signature by applying the moderated *z*-score (MODZ) procedure. We selected the HEPG2 cell line, the consensus gene signature of treated genes, and their untreated control vectors and obtained 3341 treated and corresponding 533 control conditions for 12328 genes. The *Z* difference score for each gene is then measured using the equation

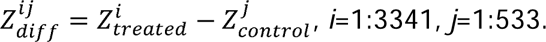

Here, 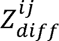 is the *Z* difference score for treated condition *i* and control condition *j*, 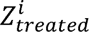 is the *Z* score of a gene corresponding to the treated condition *i*, and 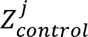 is the *Z* score of a gene corresponding to control condition *j*.

We then identified the up-and-downregulated genes for which 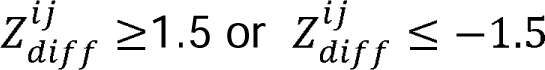 satisfies at least 60% of the control conditions, respectively.

### 2.4) Method to predict metabolic network level perturbation using the gene knockdown profile

Here, we used four inputs: (i) a generic GSMM (*iHepatocytes2322*), (ii) up-and-downregulated genes under each gene knockdown case, (iii) disease-specific gene expression data (average gene expression values under the specific disease stage), and (iv) flux state (V^ref^) of disease-specific GSMM. The knockdown-specific gene expression data were generated from the disease-specific gene expression data by considering a 2-fold up/down in the expression values of the up/downregulated genes. Next, the obtained expression data was integrated into the *iHepatocytes2322* by applying the E-Flux method, and the corresponding flux state (V^res^) was predicted by using MOMA. This process requires the flux state (V^ref^) of disease-specific GSMM. Finally, a score was assigned for each gene knockdown, similar to the method mentioned in the earlier section. We also extracted the metabolic reactions whose fluxes shifted from the diseased state toward the target state.

### 2.5) Mammalian cell culture, siRNA, and free fatty acid (FFA) treatment

Huh-7 cells were maintained in Dulbecco’s modified Eagle medium (DMEM) containing 10% fetal bovine serum (FBS) in a 5% CO_2_ incubator at 37^0^C. For siRNA gene silencing, cells were seeded at 70-80% confluency on a 6-well tissue ccellulture plate and incubated overnight at 37^0^C in 5% CO_2_. The next day 5 µM siRNA (1+2) was transfected into each well using lipofectamine 3000, following the manufacturer’s guidelines (Thermo Fisher Scientific, USA). The culture medium was changed 6 hours post-transfection with DMEM supplemented with 10% FBS, followed by incubation for 66 hours. For FFA treatment, 0.1 M stock solutions of palmitic and oleic acid were prepared in DMSO. FFA stock solution was complexed with BSA at a molar ratio of 4:1 and added to the culture medium to a final concentration of 600 and 1200 μM. FFA-treated cells were incubated for 24 or 48 hours, as indicated in the results.

### 2.6) RNA isolation, real-time quantitative PCR, western blot assay, and Nile red staining

Intracellular RNA was isolated using TRI reagent (MRC, USA), followed by reverse transcription (RT) using the FIREScript cDNA synthesis kit (Solis Biodyne, Estonia). Random hexamers were used in cDNA synthesis. SYBR green-based real-time quantitative PCR (RT-qPCR) was done as described [18]. Quantitation of GAPDH RNA level was used for normalizing the RT-qPCR data. For western blot assay, samples were resolved by SDS-PAGE and transferred to a 0.4 μm polyvinylidene difluoride (PVDF) membrane. Membranes were blocked for 1 hour at room temperature using 5% skimmed milk (in PBST). Then, membranes were incubated overnight with the primary antibody in PBST (PBS supplemented with 0.05% Tween 20) containing 5% skimmed milk at 40^0^C. Blots were washed 3 times in PBST, followed by incubation with antirabbit secondary antibody at room temperature for 1 hour. Blots were washed three times with PBST, and protein bands were visualized by enhanced chemiluminescence using a commercially available kit (Bio Rad, USA) [18]. Huh-7 cells were seeded on a coverslip for Nile red staining at 50-60% confluency. The next day, cells were transfected with BAG6 siRNA and NTsiRNA, followed by FFA (600 and 1200 μM) treatment for 24 and 48 hours, as described in the previous section. Cells were then washed twice with 1X PBS and fixed with 4% paraformaldehyde for 15 mins. The nucleus was stained with DAPI, and intracellular lipids were stained using Nile red [19].

### 2.7) Cell Viability Assay

Cell viability was measured using a commercial kit (Cell Titre 96 AQueous One Solution cell proliferation assay; Promega, Madison, USA) that uses a tetrazolium salt-based colorimetric assay [20]. 0.12*106 cells were seeded in 48 well plates and allowed to adhere overnight. The next day these cells were treated with FFA for 24 hours, the medium was removed and treated with MTT and incubated for 3-4 hours at 37^0^C in a 5% CO_2_ incubator. The absorbance was read on a multimode reader at 490 nm.

### 2.8) FACS analysis

Apoptosis was measured using Annexin V-FITC early apoptosis detection kit (Cell signalling technology, USA). Huh-7 cells were transfected with NT-siRNA or BAG6-siRNA and treated with FFA for 24 and 48 hours. Then cells were harvested by trypsin treatment and inactivated using a 10% FBS-containing culture medium. Cells were washed with ice-cold PBS. 10^5^ cells were resuspended in 1x Annexin V binding buffer. 1µl Annexin V-FITC conjugate and 12.5 µl Propidium Iodine (PI) solution were added and incubated for 10 mins in the dark on ice. The cell suspension was diluted to a final volume of 250 µl in ice-cold 1X Annexin Binding Buffer and analyzed by Fluorescence-activated Cell Sorting (FACS) [21].

## 3. Results

### 3.1) The data and preliminary results

We investigated a dataset GSE126848 [22], which includes RNA-Seq liver biopsy data of the 15 nonalcoholic fatty liver (NAFL), 16 nonalcoholic steatohepatitis (NASH), and 26 control, individuals. We obtained 5468 and 5672 differentially expressed genes (DEGs) for the NAFL and NASH categories. We also performed gene set enrichment analysis (GSEA) and found that collagen-associated pathways, apoptosis pathways, oxidative phosphorylation, DNA damage, cholesterol biosynthesis, and hypoxia-related pathways are upregulated in both groups. Collagen deposition, increased cholesterol biosynthesis, and apoptosis of liver cells are linchpins of the NAFLD progression. Hypoxia enhances cellular lipid deposition and upregulates genes involved in lipogenesis, lipid uptake, and lipid droplet formation [23]. The increase in inflammation can be observed by the upregulation of the interleukin 1 signalling and oxidative phosphorylation pathway [24,25].

Metabolic flux level profiles were predicted for each individual by integrating the gene expression data on *iHepatocytes2322* [8] (refer to Supplementary Methods). 2,859 and 1,721 reactions were altered for the NAFL and NASH categories, respectively, in which 998 were common in both categories. NAFLD is known to be strongly correlated with carbohydrates, bile acids, amino acids, and lipid metabolism dysfunction [26], which justifies the alteration of the reactions associated with these processes (**Fig. 1A-B**). The fatty acid oxidation process was downregulated for both groups, and it primarily reflects the alterations in the fatty acid activation, desaturation, and beta-oxidation pathways (**Fig. 1C** and **Data File 1**). The glycolysis or gluconeogenesis, tricarboxylic acid cycle, glyoxylate or dicarboxylate metabolism, and pyruvate metabolism were the most upregulated pathways in the carbohydrate metabolism for the NASH group. Similar results were found for the NAFL group, except some reactions were down-regulated in glycolysis or gluconeogenesis and pyruvate metabolism pathways. In the case of lipid metabolism, most reactions in the glycerolipid metabolism were upregulated, and the reactions in the arachidonic acid metabolism and glycosphingolipid biosynthesis-ganglio series were down-regulated. Most of the reactions involved in bile acid biosynthesis, recycling, and formation and hydrolysis of the cholesterol esters pathways were found to be downregulated. Significant alterations in amino acid metabolism were also noticed.

**Fig. 1:**
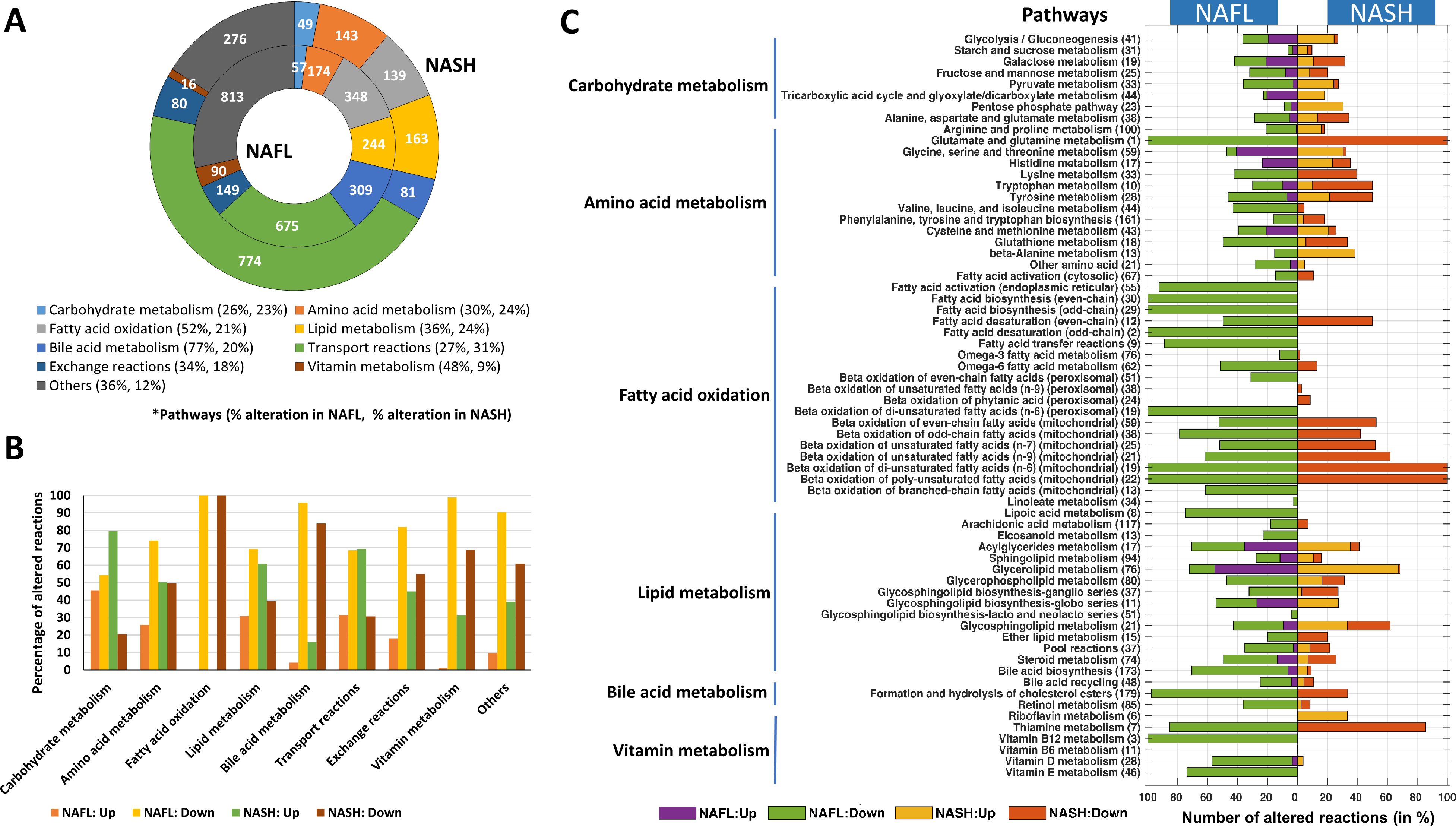
Metabolic flux level alterations. A) Represents the doughnut plot of the major metabolic processes associated with the altered reactions, and the corresponding numbers are mentioned inside. The inner and outer circles, respectively, represent the NAFL and NASH categories. Also, the level of alterations of each process is provided inside the parenthesis of the color legend. B) The bar plot represents the percentage of up-and-downregulated reactions in each metabolic process. C) Significantly altered metabolic pathways obtained for NASH and NAFL in comparison to the control group. The bar length represents the percentage of altered reactions for each pathway. Details are provided in **Data File 1**. The number inside the Y-label denotes the number of total reactions in the corresponding pathway.

### 3.2) Candidate genes capable of metabolic transformationv or reverse gene expression

The observed disease-associated alterations were further used to determine potential recovery options. At the metabolic stage, we systematically carried out a 90% gene knockdown and calculated the transformation score (TS) [15], reflecting the extent to which it may transform the disease state towards a healthy state (**Fig. 2A** and **Data File 2)**. Genes with positive TS (hereafter referred to as metabolic candidate genes, *MCG*) were selected for further study. These obtained *MCG* for both groups are enriched in retinol metabolism, one carbon pool by folate, tyrosine metabolism, drug metabolism, pyruvate metabolism, glycolysis/gluconeogenesis, and alanine, aspartate, and glutamate metabolism (**Fig. 2B-C**).

**Fig. 2:**
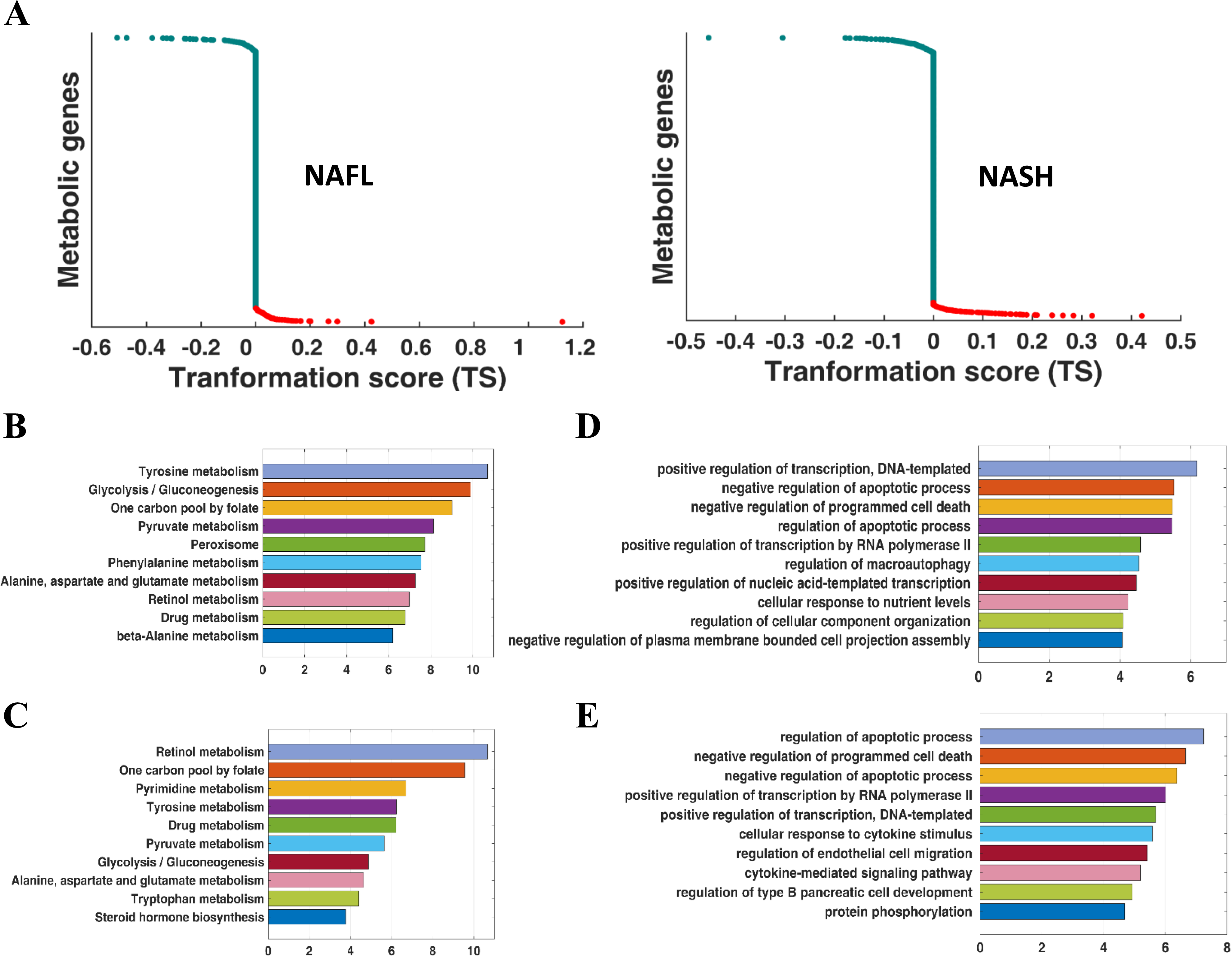
Transformation score (TS) and connectivity map (CMap) analysis. A) TS of the metabolic genes, obtained by performing 90% gene knockdown exercise. Here, the red dots represent the genes with TS>0. B-C) The top10 enriched pathways of the metabolic genes with positive TS for NAFL and NASH group, respectively. D-E) The top10 enriched pathways enriched with the candidate genes for NAFL and NASH group, respectively.

We also obtained a set of genes whose knockdown may result in a reverse gene expression profile to that of the DEGs associated with NAFLD landscape using the Connectivity Map database (CMap) [16] for the HEPG2 cell lines and termed them as candidate proteins (*Cp*) (**Data File 2)**. These obtained *Cp* for both categories are enriched in cell death-related processes (**Fig. 2D-E**).

### 3.3) PPI network analysis of *MCG*, *Cp*, and DEGs identified crucial proteins

The intricate interactions of proteins regulate the changes in both the transcriptomic and metabolic networks. We collected the DEGs, *Cp*, and *MCG* for each group and constructed the directional PPI networks (**Fig. 3A-B,** refer to Supplementary Methods). The structural controllability theory [27] was next applied to identify the minimum number of driver nodes capable of controlling the whole network. The nodes were then graded into indispensable (*I*), dispensable, and neutral categories based on the number of driver nodes. In the NAFL and NASH categories, 45% and 44% of nodes were drivers, and 18.67% and 20.59% were indispensable, respectively (**Fig. 3C**). We used the relation J_CP_ == J n Cp to find the set of indispensable candidate proteins (*ICp*) in the network, where *I* = set of indispensable proteins and *Cp* = set of candidate proteins. NAFL and NASH categories contained 15 and 29 proteins respectively as *ICp* (**Fig. 3D**). These are the most crucial proteins in the network because they generate reverse gene expression profiles and also influence network controllability [3].

**Fig. 3:**
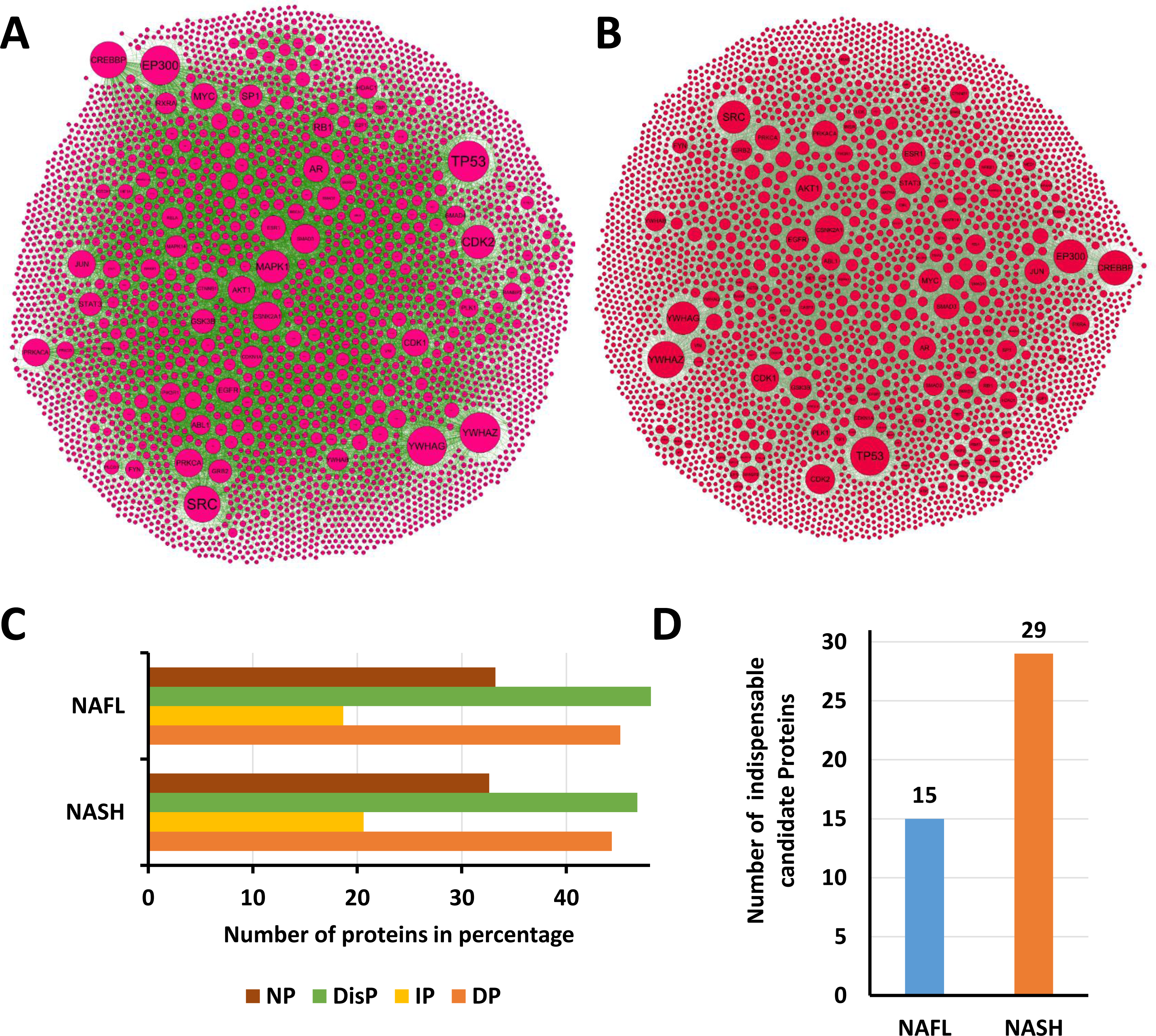
Protein-protein interaction network analysis. A-B) The PPI network of the NAFL and NASH groups, respectively. Here the greater the node size, the larger is its degree. The NAFL network consists of 3181 nodes and 16791 interactions, while the NASH network consists of 3195 nodes and 16703 edges. C) The abundance of neutral (NP), dispensable (DisP), indispensable (IP) and driver proteins (DP) nodes in the two networks. D) Number of indispensable Candidate proteins (ICp) in the NAFL and NASH group.

### 3.4) Potential therapeutic targets in NAFLD

The knockdown effect of the indispensable candidate proteins (*ICp*) on the disease-specific GSMMs was investigated to capture the metabolic flux level transition from the disease state to the target state. We calculated the TS for each *ICp* (**Fig. 4A**) and considered the proteins with positive TS as potential targets. We obtained three targets for NASH, namely, BAG6, CYCS, and CASP3. The significance of these targets in the NASH landscape was portrayed through their interactors in the NASH network. CASP3 has the maximum interactors, followed by BAG6 and CYCS (**Fig. 4B-E**). Their interactors contain proteins which play critical roles in the pathogenesis of NASH, for instance, CFLAR, TNFR1 for CASP3; TRAF6 for BAG6; and TP53 for CYCS [28–31]. These potential targets did not share a common first neighbour (**Fig. 4F**). So, there is no single drug or compound whose implementation can simultaneously affect all three targets. However, at an individual level, BAG6 and CASP3 are connected by ATN1 and CDKN1A; and CASP3 and CYCS are connected by BID and CASP9. The biological processes associated with the proteins of each network are shown in **Fig. 4G-I**. In the NAFL category, eight proteins were identified as potential targets, which are significantly associated with NAFLD (see Supplementary Information). The topological properties of the potential targets of both categories are provided in **Supplementary Table S1**.

**Fig. 4:**
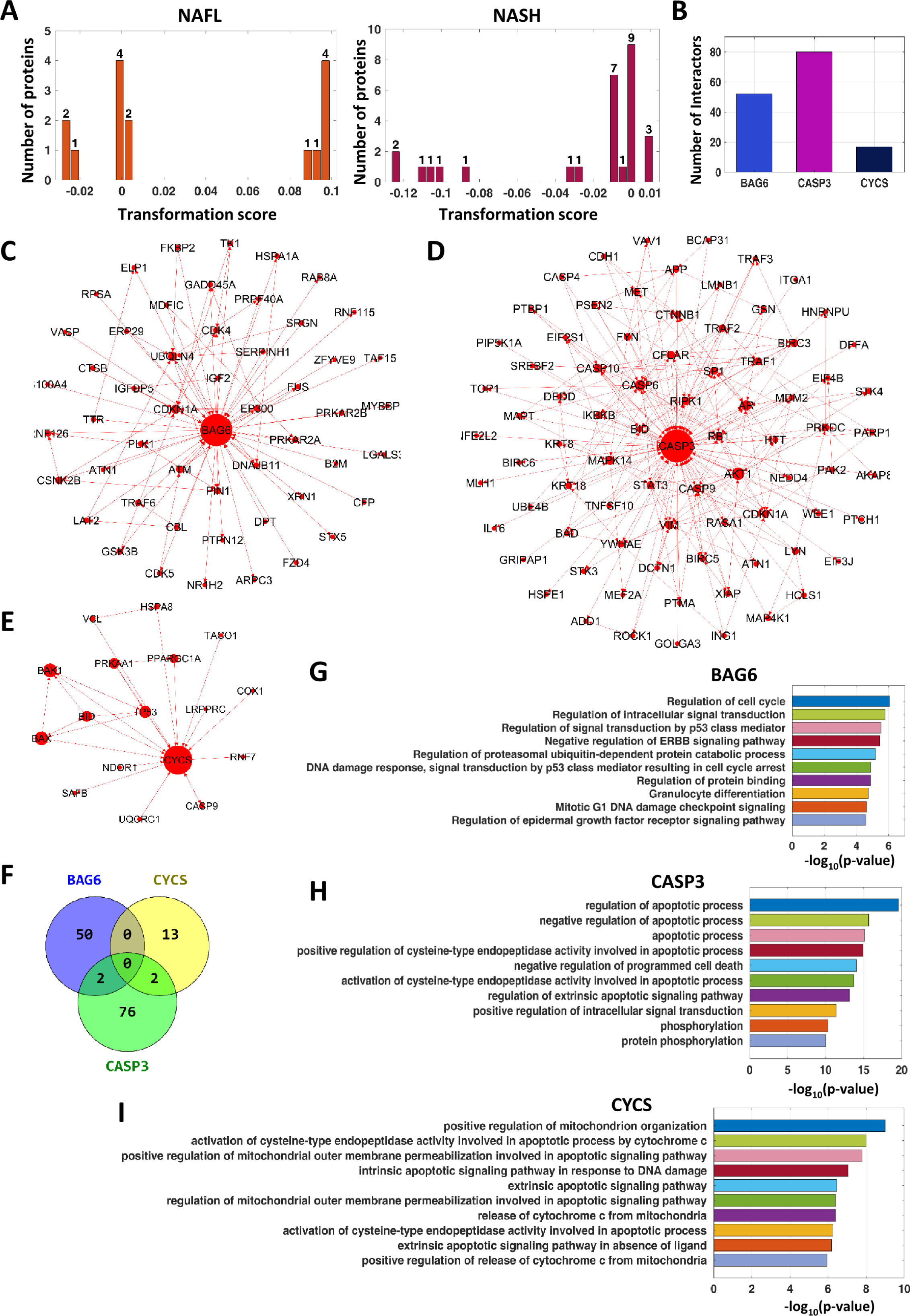
The potential targets in NAFL and NASH. A) The histogram of transformation scores (TS) per protein in ICp. The TS is plotted in the X-axis, while the bar length denotes the number of proteins. A gene is considered as a potential target if it has a positive TS. B) The targets in the NASH category and their number of interactors in the network. C) C-E) The interactors of the three targets of NASH in the network. F) Venn diagram of the interactors of the targets. G-I) The top10 enriched biological processes of interactors of BAG6, CYCS, and CASP3, respectively.

### 3.5) Capturing the effects of the potential targets on the disease-associated altered pathways

NASH is mainly driven by certain characteristics such as steatosis, hepatic cell death, inflammation, and fibrosis [32]. To comprehend the mechanistic understanding of these potential targets, we evaluated their knockdown effects on these NASH-specific characteristics (**Fig. 5A**). We captured the number of genes perturbed due to the knockdown of each target in NASH (**Fig. 5B)** and NAFL **(Fig. 5C).** The potential targets were then further investigated, keeping the focus on NASH.

**Fig. 5:**
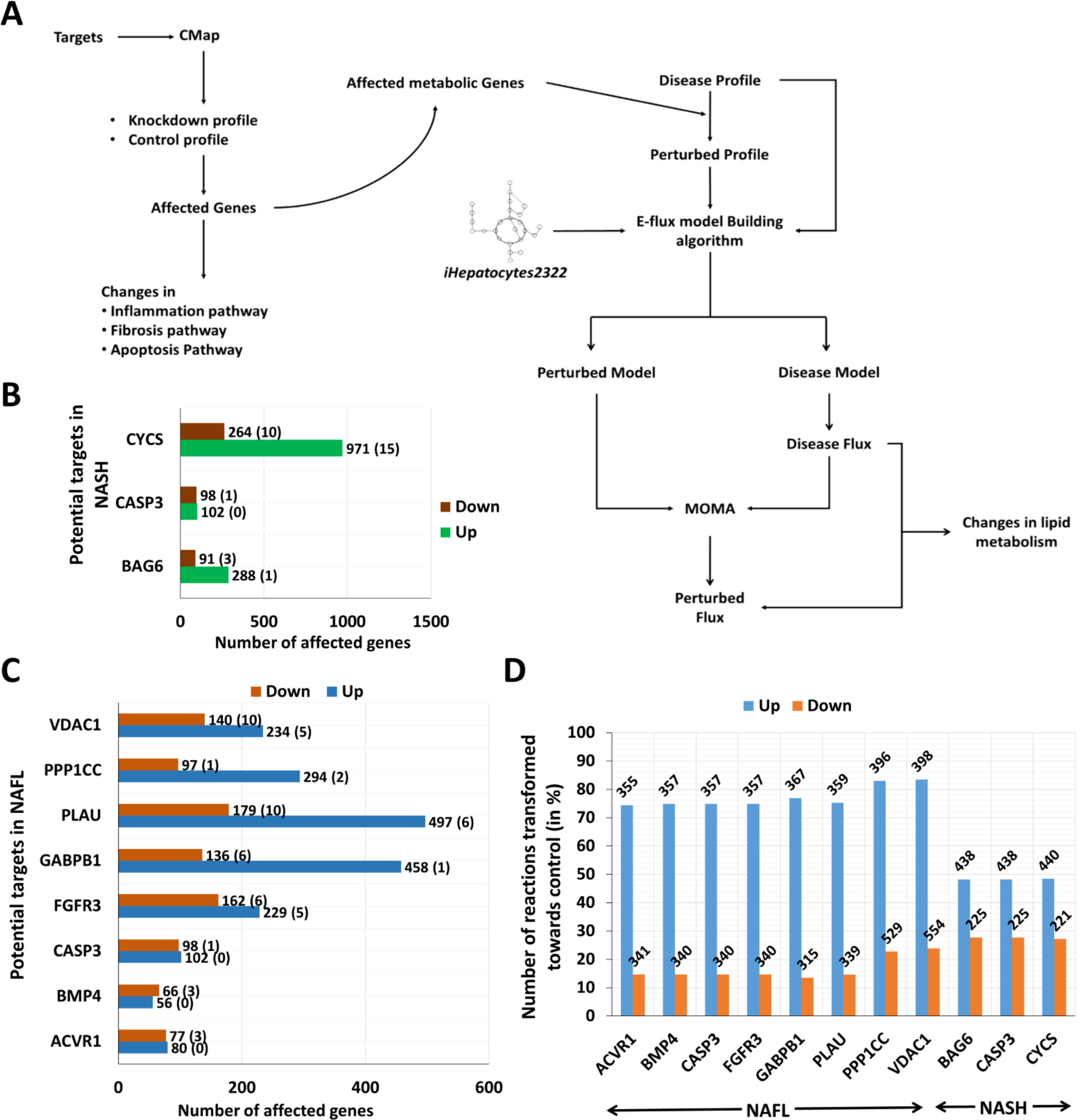
Mechanistic understanding of the potential targets. A) Schematic diagram for exploring the potential targets. The process starts with extracting the gene knockdown profile of each target from the CMap database and ends with identifying changes in the disease-specific traits such as lipid accumulation (steatosis), hepatic cell death, inflammation, and fibrosis. B-C) The bar plot of the numbers of affected genes following the knockdown of the potential targets in the NAFL and NASH groups, respectively. The total number of affected genes is shown at the top of each bar, while the number of metabolic genes is shown in brackets. D) The figure illustrates the number of reactions transferred towards normal following the knockdown of each target in their respective category, and the value is provided at the top of each bar. The number of up-and-downregulated reactions in the NAFL category are 477 and 2382. In the NASH category, these numbers are 909 and 812, respectively.

Our results revealed that the knockdown of CASP3 downregulates a fibrogenic gene, FN1, which plays a crucial role during liver fibrosis [35]. The knockdown of CASP3 can also regulate fibrosis development by upregulating WIF1, an inhibitor of the Wnt/β-catenin signalling pathway [36]. Its knockdown can regulate the inflammatory process by downregulating the inflammation inducer S100A4 [37]. Knockdown of CYCS downregulates IL32, a NAFLD-related hepatic cytokine, modulating the hepatic inflammation. Tailoring CYCS to hepatic fibrosis, we found that its knockdown downregulated Galectin 1, which can ameliorate fibrosis by inducing apoptosis to HSCs [38]. The knockdown of BAG6 upregulates MMP1 gene that can attenuate hepatic fibrosis via collagen type I and III breakdowns [39]. BAG6 knockdown downregulates the fibrogenic gene FN1 and DAG1. The latter plays a critical role in forming a basement membrane by organizing extracellular matrix proteins during liver fibrosis. It is upregulated during hepatic stellate cell activation [40]. Its knockdown also downregulates an apoptotic gene, TNFRSF21, which plays a role in hepatic cell death.

Further, to elucidate the beneficial effects of the identified potential targets on hepatic steatosis, the knockdown profile of each target in the disease-specific GSMM was integrated (refer to Method). The percentages of upregulated reactions transferred towards the control were higher than the downregulated ones for all these targets (**Fig. 5D**). All three NASH targets have a similar impact (**Fig. 6A**). The reactions involved in the fatty acid activation and mitochondrial beta-oxidation were already downregulated in NASH. The knockdowns of these potential targets were found to increase the flux rates of 66% altered reactions in the fatty acid oxidation pathways, including fatty acid activation and mitochondrial beta-oxidation (**Fig. 6B** and **Supplementary Fig. S1**). Additionally, our proposed targets can revert some of the altered carbohydrate, amino acid metabolism, glycerolipid, and sphingolipid metabolism reactions (**Data File 3** and **Data File 4)**.

**Fig. 6:**
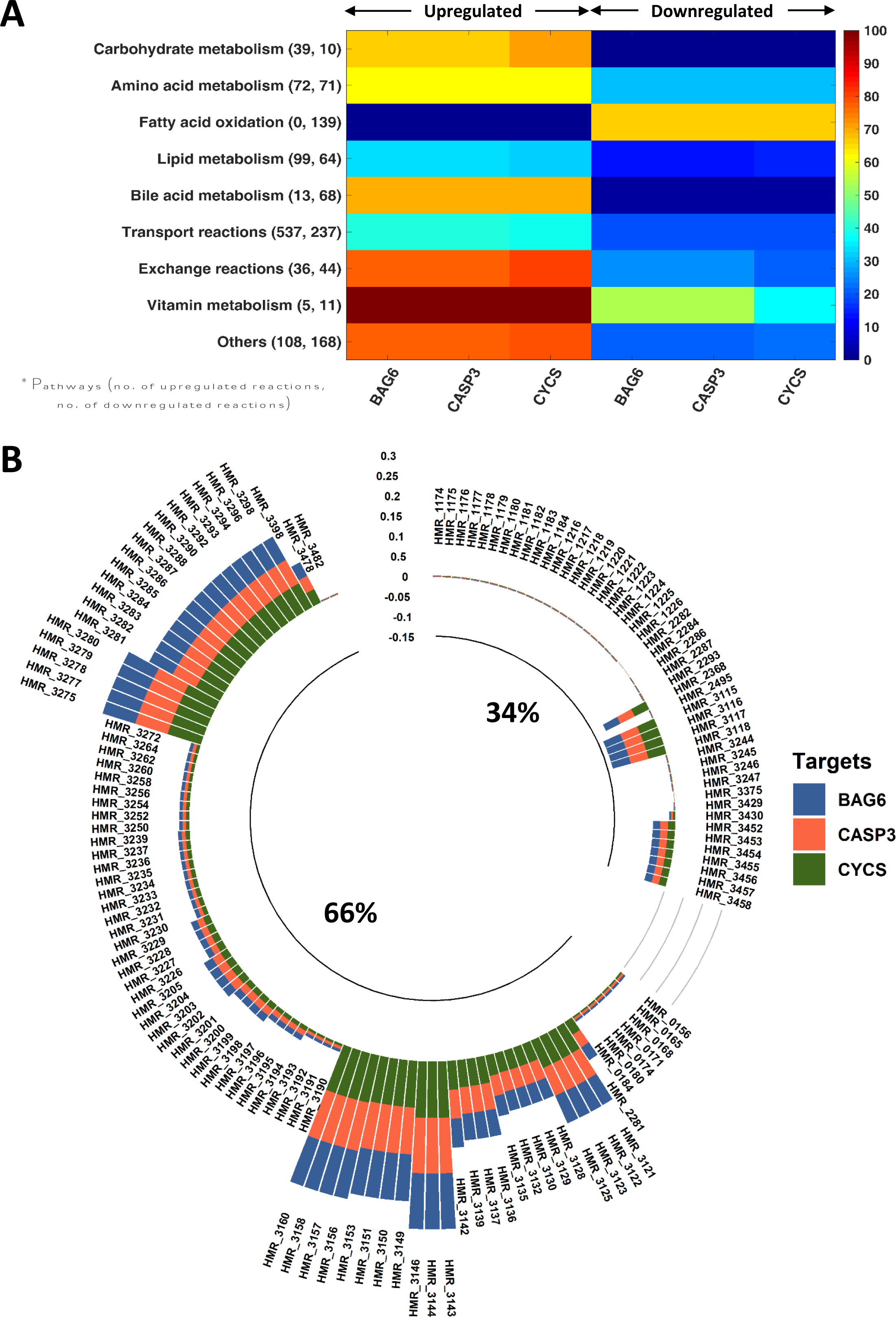
Knockdown effect of the potential targets in the metabolic level alterations. A) This figure depicts the effect of each target on the metabolic processes altered in NASH. The numbers of altered reactions transferred towards control are shown here in percentages, while the number of total altered reactions is provided inside each process’s parenthesis. Details of these reactions are provided in **Data File 3** and **Data File 4**. B) Knockdown effect of the potential targets of NASH in the downregulated reactions of fatty acid oxidation. The bar plot represents the observed flux rate differences between the knockdown and NASH conditions. A reaction is placed in two categories depending on its decreased or increased flux rate.

Thus, the knockdown profiles of these targets revealed that silencing CYCS and CAPS3 could improve inflammation, fibrosis, hepatic steatosis, and apoptotic pathways, whereas silencing BAG6 ameliorates the latter three (The proposed mechanism of these targets is summarized in **Table 1)**. Among these three targets, CYCS and CASP3 were previously found to be associated with NASH [33,34]. BAG6 is a novel target identified by our analysis. To confirm our prediction, we further evaluated the functional significance of BAG6 in the development of NASH in a cell-based model.

**Table 1:** The potential targets of NASH and their therapeutic effect.

| **Targets** | **Steatosis** | **Fibrosis** | **Inflammation** | **Cell death** |
| --- | --- | --- | --- | --- |
| **BAG6** | Increases flux rates of the reactions involved in fatty acid activation and mitochondrial beta-oxidation. | Downregulates FN1 and DAG1, and upregulates MMP1. | Not found | Downregulates TNFRSF21. |
| **CASP3** | Increases flux rates of the reactions involved in fatty acid activation and mitochondrial beta-oxidation. | Downregulates FN1 and upregulates WIF1. | Downregulates S100A4. | Inhibiting CASP3 protects against hepatocellular damage and cell death (50). |
| **CYCS** | Increases flux rates of the reactions involved in fatty acid activation and mitochondrial beta-oxidation. | Downregulates Galectin1. | Decreases expression of cytokine IL32. | CYCS release from mitochondria to the cytosol is a key process in initiating apoptosis (51). |

### 3.6) BAG6 plays a critical role in intracellular lipid accumulation and expression of genes associated with NASH

We used a human hepatoma (Huh7) cells based model of NASH to evaluate the significance of BAG6 in NASH. Huh7 cells were treated with free fatty acid (FFA) to pave the way for lipid droplet accumulation and upregulation of proteins involved in NASH, recapitulating some of the NASH-hallmarks [19]. No effect on cell viability was observed when these cells were treated with 600 μM and 1200 μM of FFA for 24 and 48 hours, respectively (**Supplementary Fig. S2A**). Nile red staining showed intracellular accumulation of fatty acids at both concentrations (**Supplementary Fig. S2B**). 48-hour treatment with FFA showed a more pronounced signal compared to its counterpart (**Supplementary Fig. S2B**). Aliquots of the same samples were used to measure the expression of NASH-signature genes such as inflammatory cytokines: interleukin-8 (IL-8), tumour necrosis factor-alpha (TNFα), fibrogenic factors dystrophin-associated glycoprotein 1 (DAG1), fibronectin 1 (FN1), fibrogenic factors alpha 2-microglobulin (α2-M), connective tissue growth factor (CTGF), and vascular endothelial growth factor (VEGF). In agreement with the earlier report [19], there was a significant increase in the transcript levels of the above-mentioned genes (**Supplementary Fig. S2C**). Next, Huh-7 cells were transfected with siRNAs against BAG6 for 72 hours, followed by measurement of the level of BAG6 protein by western blot analysis using a BAG6 antibody. Both BAG6 siRNA-1 and BAG6 siRNA-2 significantly reduced the level of BAG6 protein, and cotransfection of both siRNAs was more effective. A non-targeting siRNA was used as a control (**Fig. 7A,** upper panel). GAPDH level was monitored in aliquots of the same samples to ensure equal loading (**Fig. 7A**, lower panel). Measurement of cell viability in samples processed in parallel ruled out any cytotoxic effect of the siRNAs (**Fig. 7B**). Further, we tested the impact of BAG6-siRNA on apoptosis. No significant alteration in the percentage of apoptotic cells was observed in BAG6-siRNA transfected cells in the presence or absence of FFA (**Supplementary Fig. S3**). Next, the level of fatty acid accumulation was measured in BAG6-siRNA transfected cells. Ablation of BAG6 reduced intracellular fatty acid accumulation in the Huh7 cells treated with 1200 μM FFA for 48 hours (**Fig. 7C**). Estimating the percentage of Nile red stain-positive cells in five random fields showed an average of 30% and 5% Nile red stained cells in NT-siRNA and BAG6-siRNA transfected 1200 μM FFA-treated cells, respectively (**Fig. 7D**). Next, RT-qPCR analysis of aliquots of the above samples revealed no significant increase in the expression of NASH-signature genes, whereas 1200 μM FFA treatment significantly upregulated their level in NT-siRNA transfected cells (**Fig. 7E**). All these data support the importance of BAG6 in the development of NASH.

**Fig. 7:**
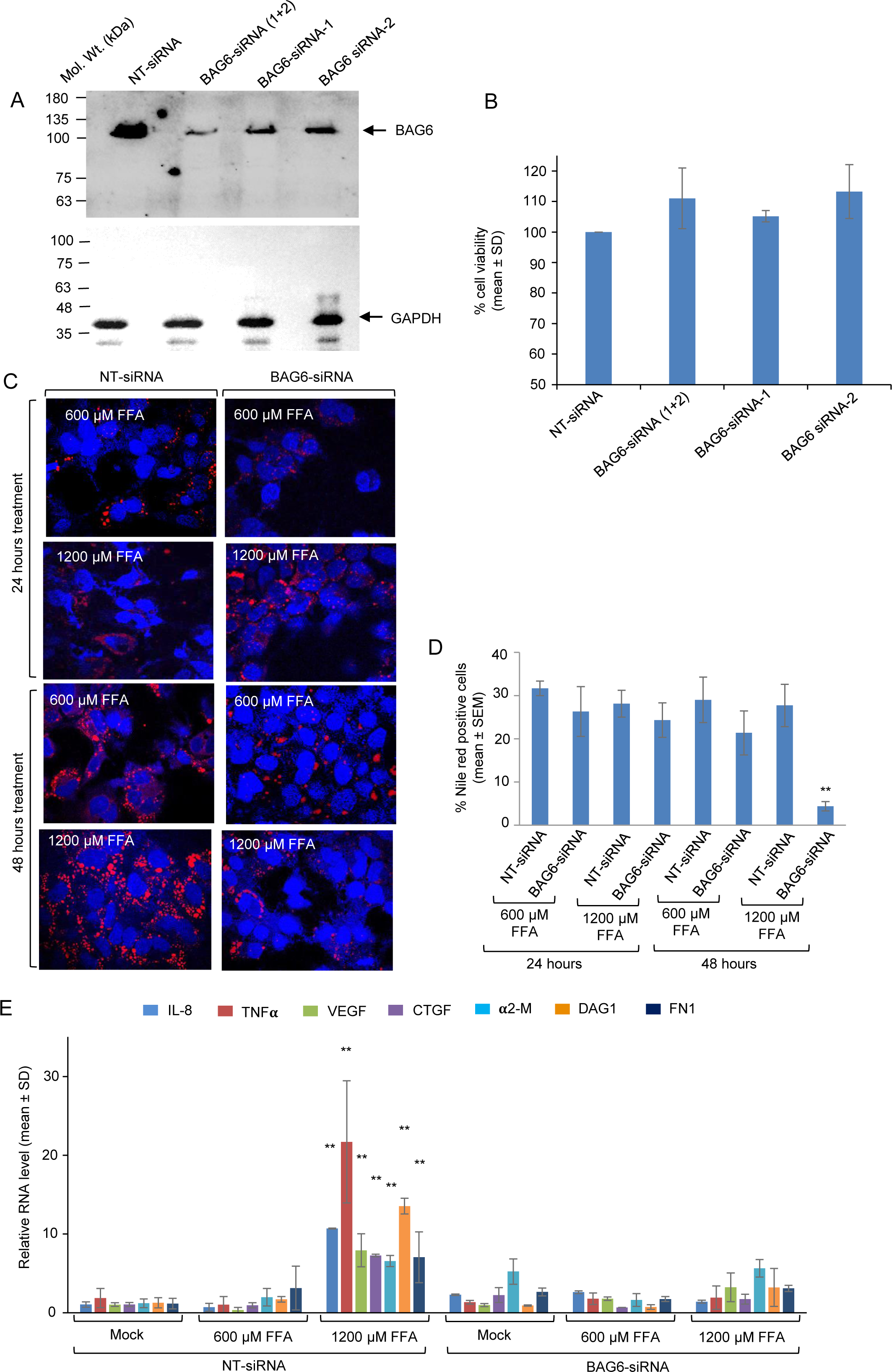
Role of BAG6 in NASH. (A) Western blot analysis of Huh7 cell lysate transfected with NT-siRNA, BAG6-siRNA(1+2), BAG6-siRNA-1, BAG6-siRNA-2, using anti-BAG6 (upper panel) and anti-GAPDH (lower panel) antibodies. (B) Percentage viability of Huh7 cells transfected with NT-siRNA, BAG6-siRNA(1+2), BAG6-siRNA-1, and BAG6-siRNA-2. Values are mean±SD of three independent experiments. (C) Immunofluorescence staining of Huh-7 cells transfected with NT-siRNA and BAG6-siRNA and treated with 600µM and 1200µM FFA for 24 and 48 hours, as indicated. Shown are merged images of nuclei (blue) and nile red-stained lipid (red). Magnification: 600X. (D) Graphical representation of mean (±SEM) of % Nile red positive cells estimated from five random fields of samples shown in (C). (E) RT-qPCR of Huh7 cells transfected with NT-siRNA and BAG6-siRNA treated with 600µM and 1200µM FFA for 24 and 48 hours, as indicated. Values are mean±SD of three independent experiments. “**” refers to P<0.01.

## 4. Discussion

NASH is a progressive multifactorial disease that initiates from a benign NAFL and may move to severe cirrhosis and liver failure [1]. Currently, it is the main reason for global liver transplants, and unabated, the numbers are going to rise only. NASH is driven by various inflammatory pathways, multiple fibrosis pathways, and metabolic alterations such as lipid metabolism, pathways associated with lipotoxicity, and antioxidant defense [1]. However, despite being studied for decades, the molecular mechanism governing NASH is yet to be deciphered, thus making the disease very hard to control. As a result, seeking potential therapeutic targets for NASH has become a top priority. Therefore, we have endeavoured to study NASH at both the gene and metabolic levels to identify potential targets with high confidence.

The collaborative study of metabolic and PPI networks opens the door to capturing the effects of disease progression in multiple layers and offers more realistic solutions to understand the disease mechanism. We first identified the significant elements of the genomic and metabolic level alterations. We obtained the candidate proteins (*Cp*) based on the former, which can initiate a reverse gene expression profile. We constructed a PPI network with nodes consisting of DEGs, *Cp*, and metabolic genes with a positive transformation score (TS) to demonstrate our hypothesis that these proteins may contain potential targets. The control theory algorithm identified the driver nodes from the network that eventually resulted in indispensable nodes, which are the most fragile in the network [2]. These nodes are prone to mutations and are often targeted by viruses and drugs [2,3]. We identified the indispensable candidate proteins (*ICp*) (the common nodes between the *Cp* and the indispensable proteins) and checked their knockdown effects on the disease-specific GSMMs. The calculated TS were then used to quantitatively determine their effectiveness. Finally, *ICp* with positive TS were deemed potential targets because they can induce reverse gene expression to the DEGs, affect the controllability of the network, and revert the disease flux state towards control.

Using our methodology, we have identified three potential targets in NASH, BAG6, CYCS, and CASP3. These three proteins could cause disease reversibility at both the protein and metabolic levels, and affect the controllability of the NASH network. Using *in-silico* gene knockdown, we have shown that these proteins are associated with fibrosis, apoptosis, inflammation, and fatty acid accumulation. Among these three targets, BAG6 was previously not reported to be associated with NASH. We were interested in evaluating our prediction on the novel target with an *in-vitro* experiment. So, we performed an *in-vitro* experiment of the Huh-7 cell line with BAG6 knockdown and monitored its effect on NASH.

BAG6 (BAT3/Scythe) is a ubiquitin-like protein involved in myriad non-related physiological and pathological processes, including apoptosis, antigen presentation, immunological pathways, and T-cell response. It is cleaved in the cytosol by CASP3 in response to intrinsic or extrinsic apoptosis, yielding a C-terminal fragment of BAG6 that induces apoptosis [41]. Again, the exosomal BAG6 activates the NK cells, while the soluble BAG6 inhibits the NK cell cytotoxicity [41]. Here, we observed that BAG6 siRNA transfected Huh-7 cells have reduced intracellular fatty acid accumulation compared to free fatty acid (FFA) treated cells, proving the beneficial effect of BAG6 inhibition on hepatic steatosis. Its impact on hepatic fibrosis development was further confirmed by RT-qPCR analysis, where we found the reduced expression of some fibrogenic factors like α2-M, CTG, DAG1, FN1, and VEGF following BAG6 knockdown. The beneficial effect of BAG6 inhibition in controlling the inflammation was further eminent because the RT-qPCR analysis revealed no significant increase in the expression of IL-8 and TNFα. In contrast, FFA treatment significantly upregulated their level in NT-siRNA transfected cells. These findings support our proposed hypothesis that the knockdown of BAG6 might revert the disease state towards the control, and it can positively affect the NASH traits. Moreover, data indicated that BAG6 was not differentially expressed in NASH. Thus it’s not involved in NASH development, rather, its inhibition can reduce the disease progression. Thus, BAG6 can potentially become a suitable drug target in NASH and need further evaluation through *in-vivo* studies.

## Conclusions

The study proposes BAG6 as a novel potential target for NASH and needs further evaluation through *in-vivo* studies. The current study provides a pragmatic computational framework for capturing their effects on disease-specific characteristics.

## Limitations

In this study, we explored liver biopsy RNA-Seq data from GEO for the differentially expressed genes and altered metabolic reactions to capture the core genes responsible for the molecular changes leading to disease reversibility. However, the paucity of causal interaction information available in the literature imposed constraints on our investigation. It is plausible that a denser causal network, replete with abundant information from the connectivity map, could further reinforce the identified target genes and the underlying mechanism. Our *in-silico* findings have been validated by a cell-based model, but a more complex animal model could potentially provide a deeper therapeutic insight into BAG6 as a target for NASH.

## Highlights

- Explore a human liver biopsy data to understand molecular alterations in NASH.
- Collaborative study of PPI and metabolic network reveals the potential targets.
- Identifying genes capable of disease reversibility.
- Elucidating the knockdown effects of the potential targets in the NASH hallmarks.
- Experimentally evaluated the functional significance of BAG6 in developing NASH.

## Ethical approval

The studies described here were carried out in accordance with the guidelines of THSTI Institutional Biosafety Committee.

## Availability of data and materials

**Data File 1, 2, 3**, and **4** can be found in the GitHub repository: https://github.com/samrat-lab/NASH_BAG6

## Author contributions

SC conceived the study. DTS, AP, and SC designed the study; DTS performed the protein-protein interaction network analysis; AP performed the metabolic network analysis; MS designed the *in vitro* experiments; UB performed the *in vitro* experiments; SC and NB supervised the work; DTS, AP, UB, MS, NB, and SC wrote the manuscript.

## Abbreviations

BAG6: BCL2 Associated Athanogene 6
NAFLD: Nonalcoholic fatty liver disease
NASH: Nonalcoholic steatohepatitis
NAFL: Nonalcoholic fatty liver
PPI: Protein-protein interaction
GSMM: Genome-scale metabolic model
CMap: Connectivity Map database
DEG: Differentially expressed genes
GSEA: Gene set enrichment analysis.
NES: Normalized enrichment scores
MOMA: Minimization of metabolic adjustment
TS: Transformation score
MTA: Metabolic transformation algorithm
MCG: Metabolic candidate genes
CP: Candidate proteins
DPN: Directional PPI network
ICP: Indispensable candidate protein
FFA: Free Fatty Acid

